# Hypothesis, analysis and synthesis: it’s all Greek to me!

**DOI:** 10.1101/561175

**Authors:** Ioannis Iliopoulos, Sophia Ananiadou, Antoine Danchin, John P. A. Ioannidis, Peter D. Katsikis, Christos A. Ouzounis, Vasilis J. Promponas

**Affiliations:** Division of Basic Sciences, School of Medicine, University of Crete, GR-71110 Heraklion, Greece; School of Computer Science, University of Manchester, Oxford Road, Manchester M13 9PL, UK; Institute of Cardiometabolism & Nutrition, Hôpital de la Pitié-Salpêtrière, 47 blvd. de l’Hôpital, F-75013 Paris, France; School of Biomedical Sciences, Li Kashing Faculty of Medicine, University of Hong Kong, 21 Sassoon Road, SAR Hong Kong, China; Meta-Research Innovation Center at Stanford (METRICS), Stanford University, 1265 Welch Rd, MSOB X306, Stanford, CA 94305, USA; Department of Immunology, Erasmus University Medical Center, 3015 CN Rotterdam, The Netherlands; Biological Computation & Process Lab (BCPL), Chemical Process & Energy Resources Institute (CPERI), Centre for Research & Technology Hellas (CERTH), PO Box 361, GR-57001 Thessalonica, Greece; Bioinformatics Research Laboratory, Department of Biological Sciences, University of Cyprus, PO Box 20537, CY-1678 Nicosia, Cyprus

## Abstract

The linguistic foundations of science and technology have relied on a range of terms many of which are borrowed from ancient languages, a known but little researched fact from a statistical perspective. Precise definitions and novel concepts are often crafted with those — frequently used — terms, yet their etymology from Greek or Latin might not always be fully appreciated. Herein, we demonstrate that frequently used terms span almost the entire PubMed^®^ database, while a handful of terms of Greek origin retrieve 80% of all entries. We argue that the etymology of those critical terms needs to be fully grasped to ensure correct use, in conjunction with other concepts. We further propose a number of terms for genomics, using prepositions that can accurately define subtle sub-disciplines of this ever-expanding field. Finally, we invite commentary by both the science community and the humanities, for possible adoption of suggested terms, not least to avoid inaccurate usage or inappropriate notions that may compromise clarity of meaning.

## Our etymological legacy

It is seldom in academic research that we, as scientists or engineers, explore or comprehend the deeper origins and subtle meanings of fundamental terms widely used in our everyday professional lives. These specialized vocabularies are primarily introduced during high school and university education to accurately describe concepts, phenomena, methodologies and techniques. At this stage, fledgling, aspiring young students usually focus on their subject matters often disregarding the linguistic dimension, as given. Later in life we get so accustomed to those terms, that it can be hard to dwell on their deeper meanings or derivatives *(1)*, <u>while their etymologies become overshadowed by intense usage</u>. As a result, even entire disciplines may be wrongly named, thus creating terminological ambiguity *(2)*.

This etymological deficit is alleviated by our linguistic backgrounds at various levels: each language contributes at different degrees to scientific and technological progress *(3).* It is widely accepted that the <u>Greek language has provided the foundations</u> of concepts to scientific research more than any other language *(4)* and, remarkably, continues to do so — as “the knowledge of names is a great part of knowledge” *(5).* We, as scientists and native speakers, carry this heavy linguistic legacy of Greek — a language with uninterrupted usage for thousands of years. Both science and technology are literally based on methods and ideas laid out and first described in Greek during the Golden Age and the Scientific Revolution of the Hellenistic period (6). Given that Greek provides the foundations of pretty much most technical, scientific, medical and philosophical terms with a rich meaning, we hereby attempt to raise a certain awareness of crucial aspects of semantics for the global community, in the age of instant information retrieval and rapid communication. Thus, we humbly urge colleagues to delve a little deeper into the etymology of standard terms of Greek origin and examine their meaning *(7).* Not only will non-Greek speakers be surprised by the richness as well as the structure of the Greek language, despite its often inept ‘naturalization’ in English or other languages, but they will most likely better understand their own thought processes and intellectual journey into their favorite subjects *(8)*.

## Universal parallels unveiled

The subject matter of scientific and technical language goes beyond science, as it <u>directly affects how our results are communicated</u> to the wider public *(9)*. Proper choice of terms and their exact meaning, often with Greek roots, deserve further explanation. Words such as ‘analysis’, ‘system’, or ‘dynamics’, may appear simple to grasp, yet they convey meanings that might have been transmuted over the past centuries to unrecognizable levels. Case in point is the word ‘machine’ which has a notion of an ‘engine’ and yet a very different meaning: ‘machine’ derives from ‘μηχαvή’ (méchané; in Doric Greek: ‘μαχαvά’) and it largely means ‘trick’. The word ‘diagnosis’ means ‘knowing through and through’, and so forth. All words ending with the suffix *-ics* were initially adjectives, not nouns: the term ‘mathematics’ is strictly speaking incomplete in ancient Greek, and was complemented by ‘philosophy’: ‘mathematical philosophy’ is a precise term, according to its original meaning^1^ *(10)*. Nouns with the same suffix are called ‘nominalized’ adjectives and also include physics, cybernetics, economics, or even the latin-derived informatics, and other -*ics*. The suffixes *-logy* and *-nomy* have been used interchangeably (e.g. ecology, astrology *versus* economy, astronomy) — why is there a ‘biology’ but not a ‘bionomy’? This series of arguments does not necessarily imply that we need to use those terms differently, but be aware of their roots and precise meaning, so that we exact control over the scientific lingua we use when we propose or invent new terms *(11)*.

Our favourite example is of course ‘analysis’: everyone is using it, few fully understand it. ‘Lysis’ means ‘breaking up’, ‘ana-’ means ‘from bottom to top’ but also ‘again/repetitively’: the subtle yet ingenious latter meaning of the term implies that if you break up something once, you might not know how it works; if you break it twice, it means that you have implicitly reconstructed it so you understand the inner workings of the system. ‘System’ itself derives from syn- and -stema: ‘-stema’ meaning a ‘base’, a ‘component that stands up’; ‘syn-’ means ‘together’, ‘in conjunction with’, i.e. ‘syn-stema’, ‘systema’ means ‘a group of parts standing up together’. Now, you can almost speak Greek: ‘Anastema’ means ‘standing upwards’, ‘stature’ (in English). The etymology of the term ‘etymology’ points to ‘etymos’ meaning ‘real’, ‘genuine’ *(11)*. In fact, we might wonder how many PubMed^®^ abstracts contain <u>our title terms: hypothesis, analysis and synthesis</u> disjointly return 11.7 million abstracts, an impressive fraction of a total of >28 million records (July 2018)^2^.

With this level of penetration into scientific vocabularies, established over the centuries by historical legacy and an indication of social status, the use of Greek is virtually unavoidable *(8, 12).* As an <u>amusing illustration of this</u>, one could not imagine reading the table of contents of a top journal and becoming excited about a breakthrough article titled “*Co-standing rebreaking of human firstin(?) coknits identifies colorbody segregation firstins(?)”.* In reality, this article does exist, and [in proper Greek!] has been published as *“Systematic analysis of human protein complexes identifies chromosome segregation proteins*”, a very interesting study indeed — no reference is provided, as this is essentially a random sample from a vast range of possible titles. Out of the ten words carefully selected by its authors in the title of this particular example, six have a Greek root. Yet, not everyone reflects on this amazing fact — including speakers of the Greek language. These words suddenly emerge as a gift from the remote past, hidden behind Latin characters and among other widely used terms *(13).*

## One sketchy analysis

To further substantiate our argument, we have carried out a systematic analysis of all PubMed^®^ entries, to explore and quantify the presence of dominant terms that originate from Greek and convey vital meaning to the majority of the scientific literature in the biomedical domain. These <u>most frequently used terms deserve special attention</u> as they impact upon the majority of the written corpus of scientific research and training. This does not necessarily mean that less frequent terms are not critical, if ambiguous or misused — e.g. paediatrics *(14)*, but that the most frequently used terms are of paramount importance *(15)*. Evidently, proper term usage to ensure accuracy and careful dissemination of scientific results are also inextricably intertwined in our mission to keep pseudo-scientific claims at bay — e.g. physics *vs.* metaphysics, psychology vs. parapsychology, and others *(16)*.

The exhaustive analysis of term frequencies across the PubMed^®^ database, the first of its kind to the best of our knowledge, clearly demonstrates the preeminence of the Greek language in scientific literature, a fact that is <u>well-known but not explicitly addressed</u> in training curricula. For example, keeping terms that appear anywhere in more than one million entries each, we obtained a list of 243 distinct words which cover the database by almost 100% [DS01]. So-called ‘stop-words’ (words that are automatically excluded from any PubMed^®^ query) are entirely represented by English terms. One would like to exclude articles (e.g. ‘the’), prepositions (e.g. ‘to’, ‘in’) and short verb forms (e.g. ‘is’, ‘was’), and retain mostly nouns and a few adjectives, to examine how many commonly used words deriving from Greek can cover (how much of) PubMed^®^. Note that no single gene or protein name appears in the list of most frequent terms *(17)*, as those are represented by frequencies one order of magnitude smaller (10K-100K times for the most frequently used ones, e.g. lysozyme, collagen, globin, etc. — not shown). Using a part-of-speech tagger^3^, we have retained only words with rich meaning corresponding to nouns (NN*), adjectives JJ*) and verb forms (VB*), a total of 172 terms [DS02] (with >1M appearances) — subtracting 20 terms with length <4 characters^4^ we are left with 152 terms [DS03]. When combined into a disjoint query, using the OR operator [DS04], these 152 terms retrieve ~27.8M entries, the majority (~97%) of all entries in the PubMed^®^ database (28,694,193 – 28 July 2018).

From this initial dataset, we have subsequently identified 15 terms with Greek origin using etymological dictionaries^5^: these terms might require a further qualification about etymology^6^. Importantly, the constructed query composed of those 15 terms — corresponding mainly to nouns^7^ [DS05] — returns more than 23,000,000 distinct PubMed^®^ entries i.e. 83% of the above mentioned 27.8M entries, or 80% of the entire database! An <u>impressive percentage</u> indeed.

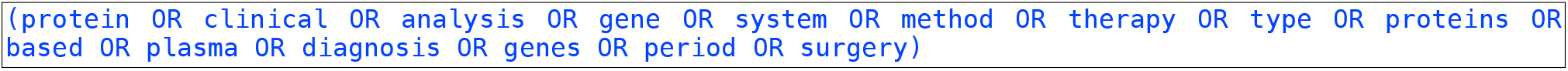

As a control, we use the 152 terms *minus* the above set of Greek words — with the NOT operator, the query being a composite of the total number of 137 frequent terms excluding the Greek ones, representing other words of different origins; note, however, that certain of these 137 query terms might be remotely connected by etymology to Greek words, for example ‘mice’/μύες. This composite query returns a mere 4,721,339 unique entries (28 July 2018) of which only 1,600,844 contain abstracts. In fact, most of the abstracts in this set also contain multiple Greek words with lower frequencies, such as ‘phenomenon’, or ‘hemorrhage’. Arguably, our estimates are on the conservative side, as low-frequency terms are not taken into account.

It should be noted that these numerical trends represent underestimates of the prevalence of Greek terms, as they do not necessarily refer to full-text documents. Another source of under-estimation, not impacting the ultimate conclusions, results from the dual presence of singular and plural forms in the query for a subset of terms; by further expanding the 15-term query to contain both forms for all terms, the reported coverage would possibly increase beyond the term expansion capabilities of Entrez^®^.

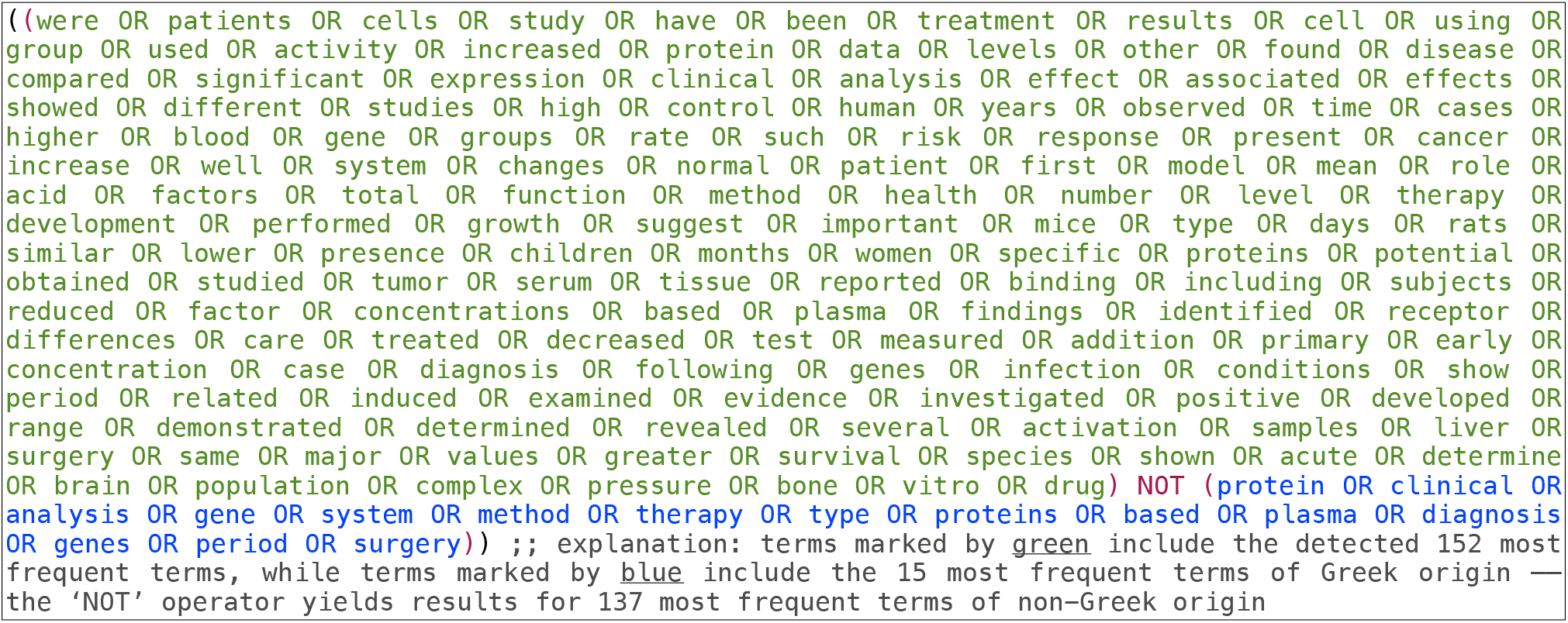

## Usage under control

The introduction of novel terms for <u>new concepts and ideas will need to be carefully crafted, following best</u> practices from the past so that our current insights do not play a role retrospectively but also prospectively *(18)*. Hence, we should strive to direct our thinking towards a highly elaborate process of inventing or introducing new terms, following other scientific fields where — by tradition — there might be more attention for precise definitions or names *(19)*. Our evolving nomenclatures should become tightly governed by greater control of language use and self-discipline. In the field of genomics, for instance, there have been creative usage patterns to qualify various genome biology sub-disciplines, for example “epigenomics”. Other – genomics / -omics terms might not be as precise, or charming — for more commentary on this particular topic, a well-known blog provides a treasure trove of such interesting definitions^8^. Case in point is the term “metagenomics”, which although difficult to revoke at this point, does not arguably carry the proper semantics with the *meta* – prefix (*meta*: meaning beyond). A better division of terms in this area might have been *endo-genomics* for host microbiomes *(endo:* meaning inside) and exo-genomics *(exo:* meaning outside) for environmental microbiomes (strictly speaking, these prefixes are not prepositions — see below). A similar situation might have occurred for the quest for life in the universe: the term *exo*-biology was gradually replaced by astro-biology, perhaps with limited linguistic (and possibly scientific) considerations (20) — one’s impression is that it had to rhyme with astro-nauts, and not exo-nauts. We conclude here with a classic recommendation: “Πάσαv γλώσσα βασάvιζε” *(tr.* “try every art of tongue” – Aristophanes, 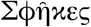 Sphekes; Latin: Vespae, 422 BC).

## A modest proposal

Evidently, all of research, training, education and knowledge transfer should take into account a much deeper appreciation and understanding of this linguistic treasure^9^, in order to <u>satisfy the needs of future conceptual advances in science</u>. For instance, a concept represented by the suffix ‘-some’ (from ‘soma’), meaning ‘body’ in Greek has already been used for proposed neologisms in the literature during recent times, capturing notions from molecular and cell biology to biochemistry and physiology (**Table 1**): while some of those have been successful, others have not. This situation emphasizes the urge to craft successful terminology as close to the conceptual roots as possible, to maintain accuracy, validity and perhaps community acceptance. One of us (AD) has proposed various modifications of existing terms ending in ‘-some’ with Greek roots that reflect concepts more accurately and even elegantly, for instance ‘schizosome’ instead of ‘divisome’, ‘coptosome’ instead of ‘spliceosome’ and ‘anxiosome’ instead of ‘stressosome’ (21).

**Table 1.**
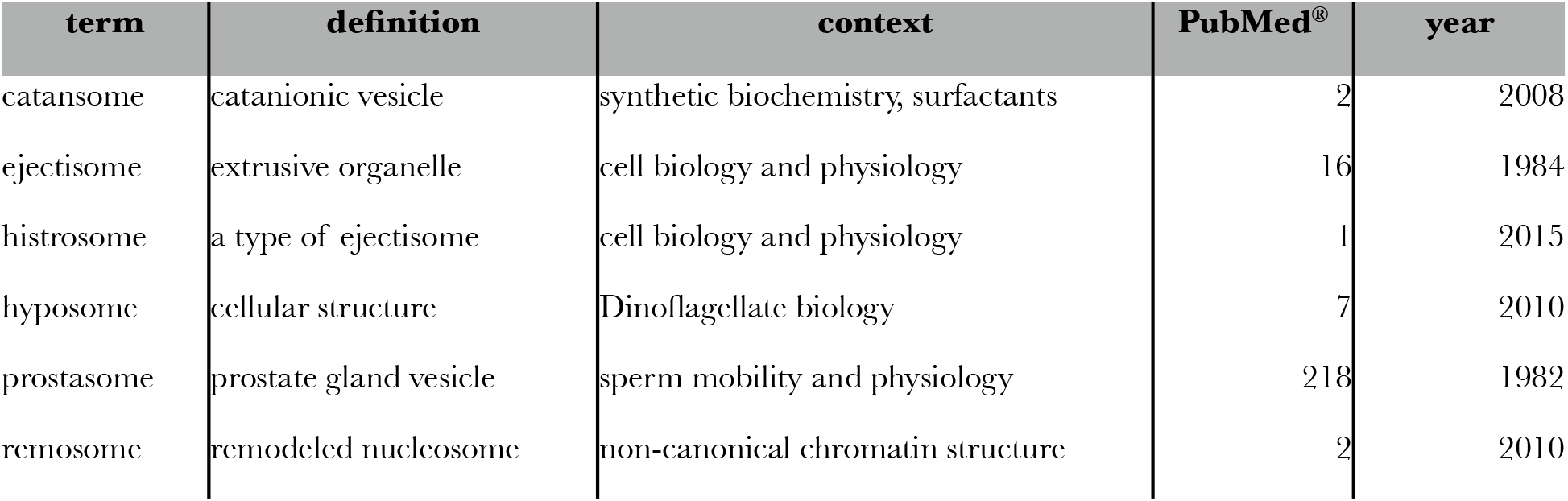
Terms that appear in the literature from different disciplines, ending in -some. Terms with suffix -some, (column one) with a short definition (column two) and the general context in the literature (column three); number of PubMed^®^ entries (column four) and year of first appearance in PubMed^®^ (column five) are also listed.

Another group of overloaded terms in modern science has been, as we have already alluded to, those with the Greek suffix ‘-ome’ (from ‘-oma’), much used and abused in recent times — and relevant to our own academic activities. To exemplify the linguistic richness of the Greek language in this specific context, we thereby explore the use of Greek prepositions to qualify and accurately specify various aspects of genomics, with their respective subject matter (**Table 2**). Our intention is not only to introduce new concepts as an exercise, but seek a more precise definition of existing activities, currently undesignated. The usage of prepositions in general conveys additional, precise and logically consistent meaning, as in the case of *epi* – genomics. Other uses, not previously proposed, are herein offered for adoption by the global community: these terms can be explored as possible means to provide novel definitions with an accurate semantic representation of corresponding, ongoing research activities. This modest proposal will hopefully <u>prevent the unfortunate use of alternative terms</u>, which might not always articulate the deeper semantics of these subfields in the genomics domain. Once more, whether frequent or not, Greek terms might come to the rescue and act as safeguards, to ensure that we are able to express subtleties of evolving scientific concepts, without retreating to artificial or untested linguistic constructs.

**Table 2.**
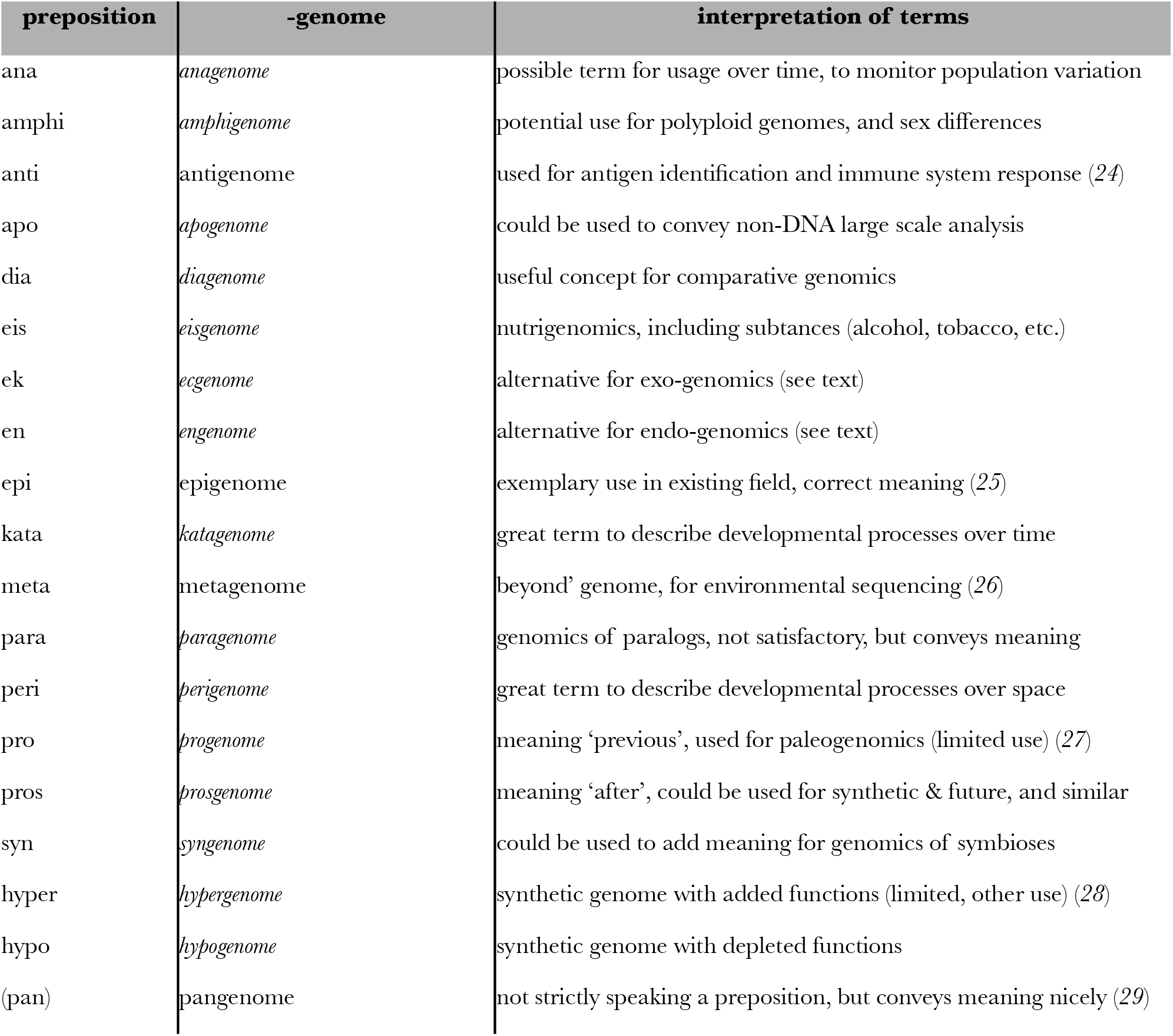
Potential term definitions using prepositions and the suffix -genome. Prepositions of Greek origin (column one) that can be associated with the suffix -genome (currently undesignated in italics, column two); interpretation of terms (column three) defines the suggested use. It can be seen that certain prepositional prefixes have been successfully used in genomics, while others have not. Certain combinations offer extremely interesting opportunities to summarize vast swathes of genome biology, qualifying specific sub-fields or aspects of the particular area of research (23).

## CONCLUSION

A number of historical factors over the past few centuries have kept apart the Greek-speaking world and turned the Greek language into a challenge, seen as ‘difficult’, maybe ‘foreign’ and definitely not very useful *(12)*. It is often forgotten that some of the greatest achievements of science, engineering, literature, philosophy, arts and architecture, were communicated in Greek, not only in the remote past but throughout the recent centuries: the Greek language greatly contributed to the development of the Renaissance, the French and American Revolutions, and modern scientific exploration. Scholars of the Western world spoke and wrote in Greek, from Newton and Leibniz to Goethe and Wittgenstein. The same situation holds for scholars who influenced the development of the life sciences, as exemplified by taxonomic names, a tradition that has survived alongside Latin since Linnaeus and Darwin. Perhaps today we can all learn a thing or two from them, now that we have the capacity to extract meaning and sense in our electronic universe with automated translation, dictionary and etymology tools, in order to achieve a precise scientific discourse and maintain accuracy of our ‘logos’ (as in -logy).

### Epi-logos: Фοβού τοβούς Δαυαούς

The proper usage of terms and composites requires careful assessment, open dialogue, scientific reasoning and deep knowledge of each specific natural science domain, as well as constructive feedback from linguists and scholars from the humanities *(8, 22)*. As an opportunity to bridge the sciences, from natural to social, we hope that this contribution will stimulate the fundamental discourse requisite to elicit consistent and accurate terminology usage in modern science and technology.

## ACKNOWLEDGMENTS

We thank Ben Blencowe (Univ. of Toronto), Georgios Floros (Univ. of Cyprus), Peter Karp (SRI International), Nikos Kyrpides (JGI Berkeley), Nikolas Papanikolaou (Univ. of Crete), Nikos Sarantakos (European Parliament) and Spyros Sfenthourakis (Univ. of Cyprus) for insightful comments and constructive criticisms.

## SUPPLEMENTARY INFORMATION

► Data supplement (and invited commentary): https://figshare.com/s733fbde421cdib5cc0245
►Etymology of the select terms of Greek origin: http://troodos.biol.ucy.ac.cy/Etymology.html

## Footnotes

1 ‘μαvθάvω’: ‘learn’; mathematics: learnable, not logical or natural philosophy; philosophy: love for wisdom.

2 https://www.ncbi.nlm.nih.gov/pubmed/?term=(hypothesis+or+analysis+or+synthesis) [accessed 28 July 2018]

3 http://cst.dk/tools/index.php

4 Most of these terms are single-letter codes, symbols (e.g. lt), abbreviations (e.g. DNA), and a few short English words (e.g. is, are).

5 https://www.etymonline.com/ (accessed, July 2018)

6 It should be noted that there is a limit to which etymology can be traced back to roots, and at times, it can be erroneous: while some decisions about etymology may cause slight variations in statistical estimates, the overall trends remain the same.

7 In fact, 13 out of the 15 terms of Greek origin are nouns.

8 phylogenomics.blogspot.com/search/label/badomics

9 ‘treasure’ arising from: *thesaurus* «θηϭαυϱός

